## Supplemental figures and legends for "Phosphoproteomics reveals rewiring of the insulin signaling network and multi-nodal defects in insulin resistance"

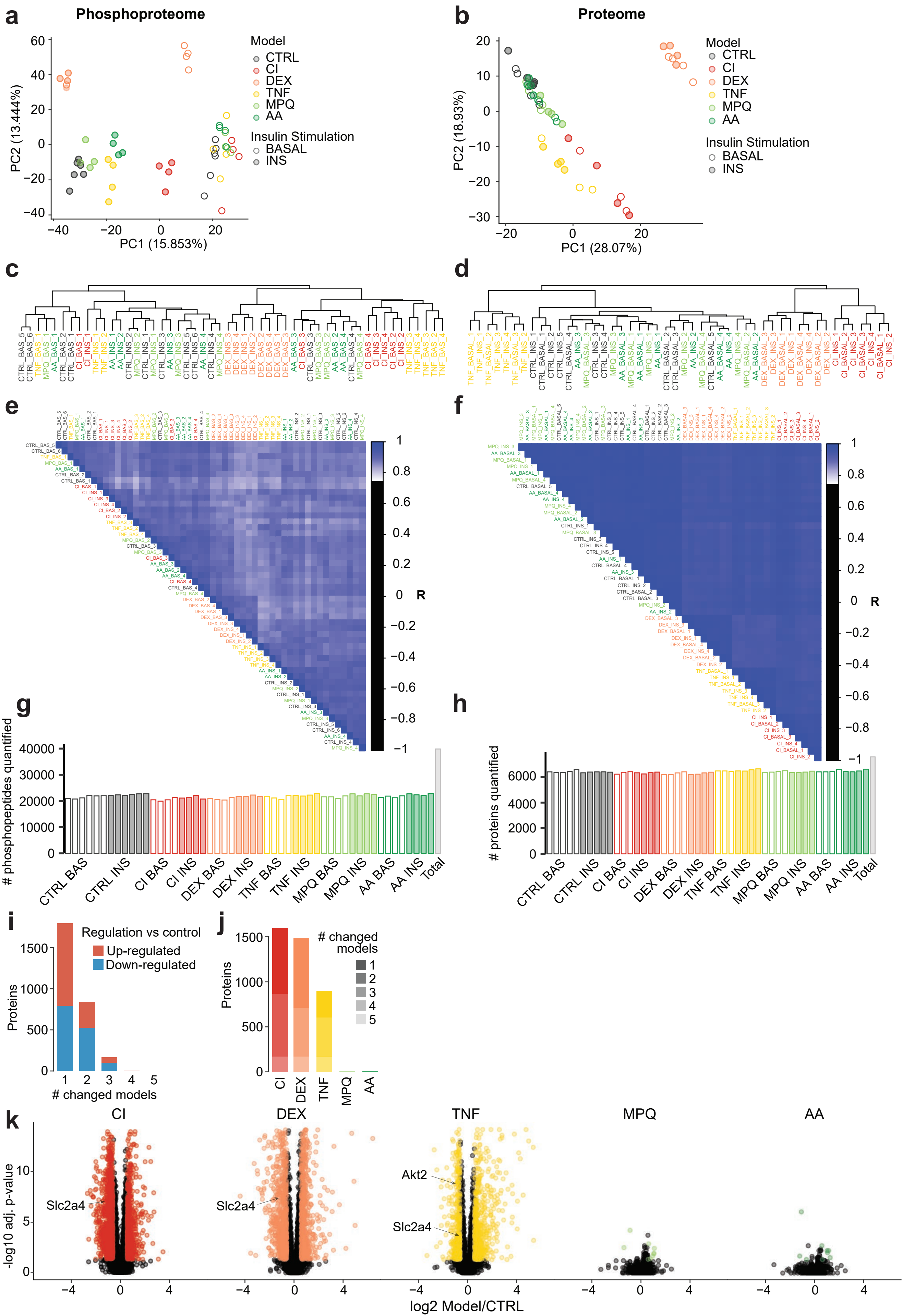

a
Phosphoproteome
PC1 (15.853%)
PC2 (13.444%)
Model
CTRL
CI
DEX
TNF
MPQ
AA
Insulin Stimulation
BASAL
INS
b
Proteome
PC1 (28.07%)
PC2 (18.93%)
Model
CTRL
CI
DEX
TNF
MPQ
AA
Insulin Stimulation
BASAL
INS
c
d
e
R
f
R
g
### phosphopeptides quantified
CTRL BAS
CTRL INS
CI BAS
CI INS
DEX BAS
DEX INS
TNF BAS
TNF INS
MPQ BAS
MPQ INS
AA BAS
AA INS
Total
h
### proteins quantified
CTRL BAS
CTRL INS
CI BAS
CI INS
DEX BAS
DEX INS
TNF BAS
TNF INS
MPQ BAS
MPQ INS
AA BAS
AA INS
Total
i
Regulation vs control
Up-regulated
Down-regulated
Proteins
### changed models
j
### changed models
Proteins
CI
DEX
TNF
MPQ
AA
k
CI
DEX
TNF
MPQ
AA
Slc2a4
Slc2a4
Akt2
Slc2a4
-log10 adj. p-value
log2 Model/CTRL

**Fig S1: Analysis of the proteome and phosphoproteome in insulin resistant adipocytes (related to Fig 1)**

(**A-B**) PCA was performed on the 3T3-L1 insulin resistance (**A**) phosphoproteome or (**B**) proteome using the base R "prcomp" function. The first two principal components (PC1 and PC2) are plotted for each replicate, and the percentage contribution of these principal components to total variance is indicated. (**C-D**) Hierarchical clustering of the (C) phosphoproteome or (D) proteome using the base R functions "dist" and "hclust". (**E-F**) Pearson's correlation between (E) phosphoproteome or (F) proteome replicates. (**G-H**) Number of quantified phosphopeptides/proteins in each replicate of the (G) phosphoproteome or (H) proteome. (**I-J**) Proteins changed up- or down-regulated in insulin resistant cells compared to control cells. (**I**) x-axis indicates the maximum number of models in which each protein is changed. (**J**) Transparency is altered to show whether proteins are only changed in the indicated model (1 model, least transparent) or are also changed in the same direction in other models (2, 3, 4, 5 models, increasing transparency). (**K**) Volcano plots showing proteomic changes in each insulin resistance model compared to control cells. Significantly altered proteins are coloured, and select proteins are labeled.

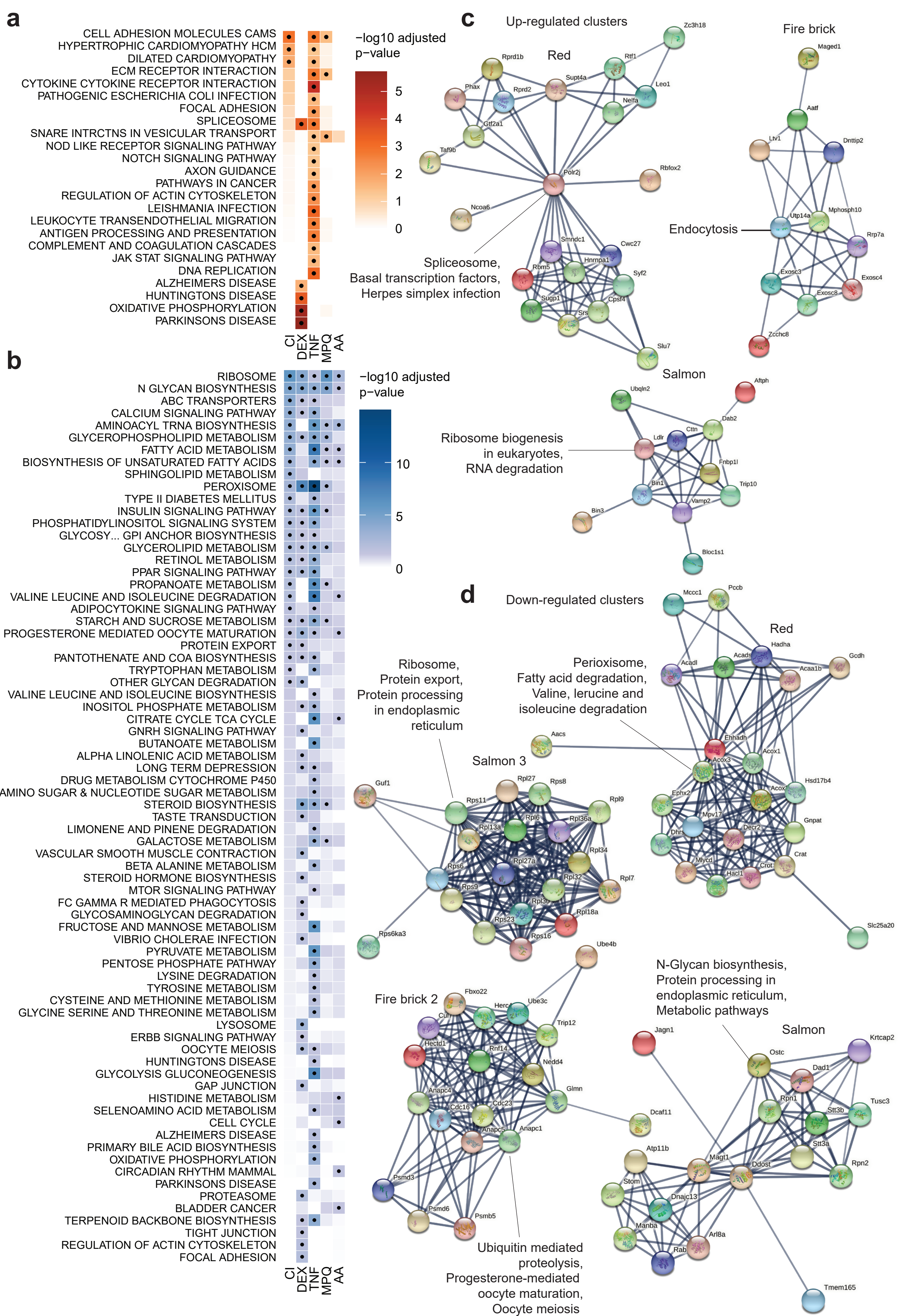
a
CELL ADHESION MOLECULES CAMS
HYPERTROPHIC CARDIOMYOPATHY HCM
DILATED CARDIOMYOPATHY
ECM RECEPTOR INTERACTION
CYTOKINE CYTOKINE RECEPTOR INTERACTION
PATHOGENIC ESCHERICHIA COLI INFECTION
FOCAL ADHESION
SPLICEOSOME
SNARE INTRCTNS IN VESICULAR TRANSPORT
NOD LIKE RECEPTOR SIGNALING PATHWAY
NOTCH SIGNALING PATHWAY
AXON GUIDANCE
PATHWAYS IN CANCER
REGULATION OF ACTIN CYTOSKELETON
LEISHMANIA INFECTION
LEUKOCYTE TRANSENDOTHELIAL MIGRATION
ANTIGEN PROCESSING AND PRESENTATION
COMPLEMENT AND COAGULATION CASCADES
JAK STAT SIGNALING PATHWAY
DNA REPLICATION
ALZHEIMERS DISEASE
HUNTINGTONS DISEASE
OXIDATIVE PHOSPHORYLATION
PARKINSONS DISEASE
CI
DEX
TNF
MPQ
AA
−log10 adjusted p−value
5
4
3
2
1
0
b
RIBOSOME
N GLYCAN BIOSYNTHESIS
ABC TRANSPORTERS
CALCIUM SIGNALING PATHWAY
AMINOACYL TRNA BIOSYNTHESIS
GLYCEROPHOSPHOLIPID METABOLISM
FATTY ACID METABOLISM
BIOSYNTHESIS OF UNSATURATED FATTY ACIDS
SPHINGOLIPID METABOLISM
PEROXISOME
TYPE II DIABETES MELLITUS
INSULIN SIGNALING PATHWAY
PHOSPHATIDYLINOSITOL SIGNALING SYSTEM
GLYCOSY... GPI ANCHOR BIOSYNTHESIS
GLYCEROLIPID METABOLISM
RETINOL METABOLISM
PPAR SIGNALING PATHWAY
PROPANOATE METABOLISM
VALINE LEUCINE AND ISOLEUCINE DEGRADATION
ADIPOCYTOKINE SIGNALING PATHWAY
STARCH AND SUCROSE METABOLISM
PROGESTERONE MEDIATED OOCYTE MATURATION
PROTEIN EXPORT
PANTOTHENATE AND COA BIOSYNTHESIS
TRYPTOPHAN METABOLISM
OTHER GLYCAN DEGRADATION
VALINE LEUCINE AND ISOLEUCINE BIOSYNTHESIS
INOSITOL PHOSPHATE METABOLISM
CITRATE CYCLE TCA CYCLE
GNRH SIGNALING PATHWAY
BUTANOATE METABOLISM
ALPHA LINOLENIC ACID METABOLISM
LONG TERM DEPRESSION
DRUG METABOLISM CYTOCHROME P450
AMINO SUGAR & NUCLEOTIDE SUGAR METABOLISM
STEROID BIOSYNTHESIS
TASTE TRANSDUCTION
LIMONENE AND PINENE DEGRADATION
GALACTOSE METABOLISM
VASCULAR SMOOTH MUSCLE CONTRACTION
BETA ALANINE METABOLISM
STEROID HORMONE BIOSYNTHESIS
MTOR SIGNALING PATHWAY
FC GAMMA R MEDIATED PHAGOCYTOSIS
GLYCOSAMINOGLYCAN DEGRADATION
FRUCTOSE AND MANNOSE METABOLISM
VIBRIO CHOLERAE INFECTION
PYRUVATE METABOLISM
PENTOSE PHOSPHATE PATHWAY
LYSINE DEGRADATION
TYROSINE METABOLISM
CYSTEINE AND METHIONINE METABOLISM
GLYCINE SERINE AND THREONINE METABOLISM
LYSOSOME
ERBB SIGNALING PATHWAY
OOCYTE MEIOSIS
HUNTINGTONS DISEASE
GLYCOLYSIS GLUCONEOGENESIS
GAP JUNCTION
HISTIDINE METABOLISM
SELENOAMINO ACID METABOLISM
CELL CYCLE
ALZHEIMERS DISEASE
PRIMARY BILE ACID BIOSYNTHESIS
OXIDATIVE PHOSPHORYLATION
CIRCADIAN RHYTHM MAMMAL
PARKINSONS DISEASE
PROTEASOME
BLADDER CANCER
TERPENOID BACKBONE BIOSYNTHESIS
TIGHT JUNCTION
REGULATION OF ACTIN CYTOSKELETON
FOCAL ADHESION
CI
DEX
TNF
MPQ
AA
−log10 adjusted p−value
10
5
0
c
Up-regulated clusters
Red
Spliceosome,
Basal transcription factors,
Herpes simplex infection
Fire brick
Endocytosis
Salmon
Ribosome biogenesis
in eukaryotes,
RNA degradation
Zc3h18
Rprd1b
Rtf1
Phax
Rprd2
Supt4a
Leo1
Nelfa
Gtf2a1
Taf9b
Polr2j
Rbfox2
Ncoa6
Smndc1
Cwc27
Hnrnpa1
Rbm5
Syf2
Sugp1
Cpsf4
Srs
Slu7
Maged1
Aatf
Ltv1
Dnttip2
Utp14a
Mphosph10
Rrp7a
Exosc3
Exosc4
Exosc8
Zcchc8
Ubqln2
Aftph
Dab2
Ldlr
Cttn
Fnbp1l
Trip10
Bin1
Vamp2
Bin3
Bloc1s1
d
Down-regulated clusters
Red
Perioxisome,
Fatty acid degradation,
Valine, lerucine and
isoleucine degradation
Ribosome,
Protein export,
Protein processing
in endoplasmic
reticulum
Salmon 3
Fire brick 2
Ubiquitin mediated
proteolysis,
Progesterone-mediated
oocyte maturation,
Oocyte meiosis
N-Glycan biosynthesis,
Protein processing in
endoplasmic reticulum,
Metabolic pathways
Salmon
Mccc1
Pccb
Hadha
Gcdh
Acadl
Acads
Acaa1b
Aacs
Ehhadh
Acox1
Acox3
Hsd17b4
Ephx2
Acox
Mpv17
Gnpat
Dhrs
Decr2
Mlycd
Crat
Crot
Hacl1
Slc25a20
Guf1
Rpl27
Rps8
Rps11
Rpl9
Rpl6
Rpl36a
Rpl13a
Rpl34
Rps6
Rpl27a
Rpl32
Rpl7
Rps9
Rpl30
Rpl18a
Rps23
Rps16
Rps6ka3
Ube4b
Fbxo22
Herc4
Ube3c
Cul1
Trip12
Hectd1
Rnf14
Nedd4
Anapc4
Glmn
Cdc16
Cdc23
Anapc5
Anapc1
Psmd3
Psmd6
Psmb5
Jagn1
Krtcap2
Ostc
Dad1
Tusc3
Rpn1
Stt3b
Stt3a
Dcaf11
Atp11b
Magt1
Ddost
Rpn2
Stom
Dnajc13
Manba
Arl8a
Rab
Tmem165

**Fig S2: Pathway and network analysis implicates functional protein modules in insulin resistance (related to Fig 1)**

(**A-B**) KEGG pathway enrichment was performed in each model by gene set test using log2 Model/Control fold change values and the KEGG pathways. (A) Pathways that are up-regulated relative to control cells. (B) Pathways that are down-regulated relative to control cells. Dots indicate significantly regulated pathways (Benjamini-Hochberg-adjusted p-value $< 0.05$). (**C-D**) Markov clustering was performed on the STRING functional networks of (C) proteins up-regulated or (D) proteins down-regulated in two or more insulin resistance models relative to control cells. All clusters containing at least ten proteins are shown, and the top 3 KEGG pathways significantly enriched in each cluster's genes are indicated. Clusters names (for example Red, Fire brick) are indicated.

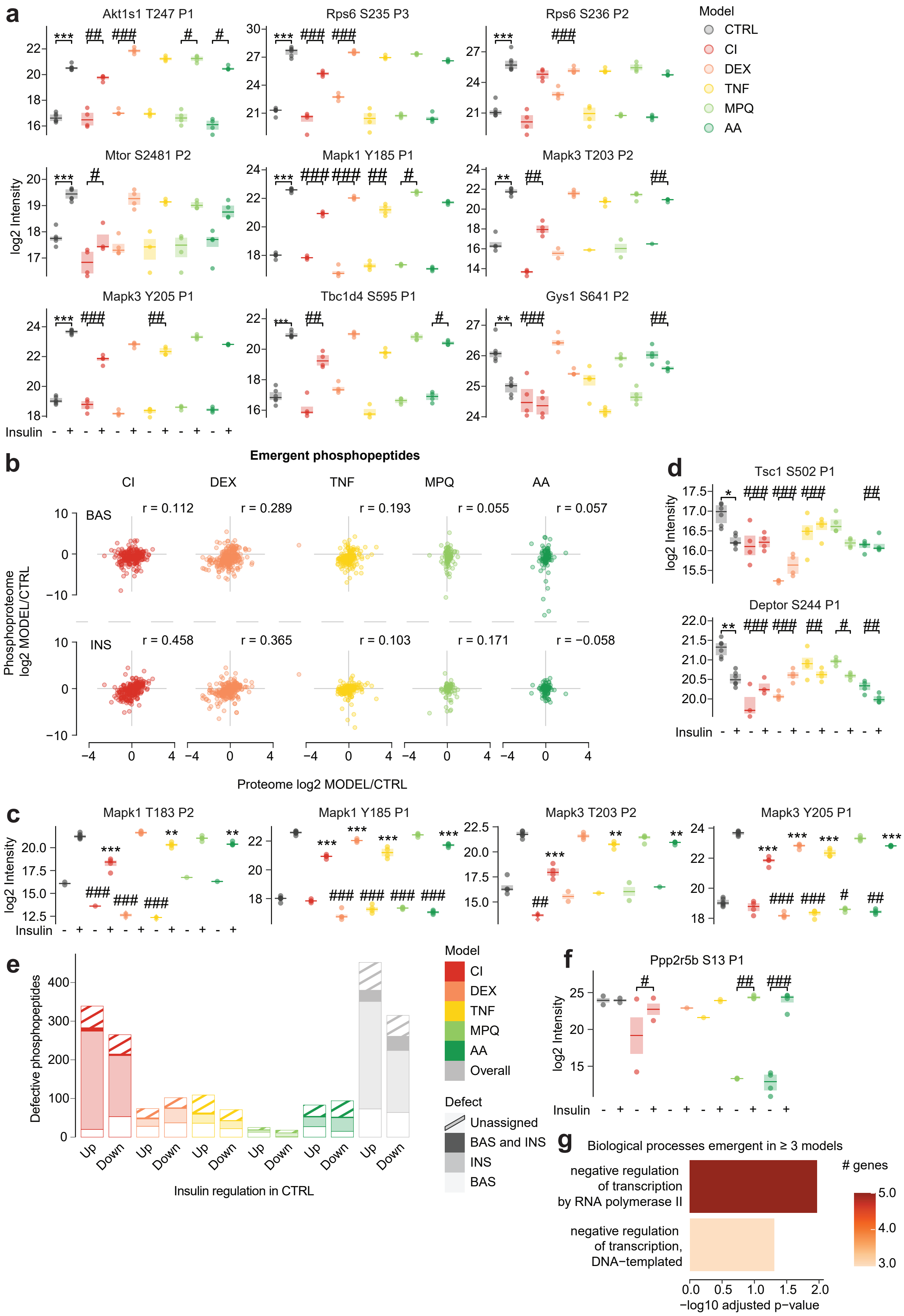

a
Akt1s1 T247 P1
Rps6 S235 P3
Rps6 S236 P2
Model
CTRL
CI
DEX
TNF
MPQ
AA
Mtor S2481 P2
Mapk1 Y185 P1
Mapk3 T203 P2
Mapk3 Y205 P1
Tbc1d4 S595 P1
Gys1 S641 P2
log2 Intensity
Insulin
b
Emergent phosphopeptides
CI
DEX
TNF
MPQ
AA
BAS
INS
r = 0.112
r = 0.289
r = 0.193
r = 0.055
r = 0.057
r = 0.458
r = 0.365
r = 0.103
r = 0.171
r = −0.058
Phosphoproteome log2 MODEL/CTRL
Proteome log2 MODEL/CTRL
c
Mapk1 T183 P2
Mapk1 Y185 P1
Mapk3 T203 P2
Mapk3 Y205 P1
d
Tsc1 S502 P1
Deptor S244 P1
e
Defective phosphopeptides
Insulin regulation in CTRL
Up
Down
Model
CI
DEX
TNF
MPQ
AA
Overall
Defect
Unassigned
BAS and INS
INS
BAS
f
Ppp2r5b S13 P1
g
Biological processes emergent in ≥ 3 models
negative regulation of transcription by RNA polymerase II
negative regulation of transcription, DNA−templated
### genes
−log10 adjusted p−value

**Fig S3: Canonical, defective, and emergent insulin signaling (related to Fig 2)**

(**A**) Select canonical insulin-regulated phosphosites. * with bracket: CTRL INS is significantly different to CTRL BAS. # with bracket: Model insulin response is significantly different to CTRL insulin response. “P1/P2/P3” indicates the number of phosphosites on each phosphopeptide. (**B**) Correlation of proteome and unstimulated (BAS) or insulin-stimulated (INS) phosphoproteome changes in phosphopeptides emergent in the indicated models. r = pearson’s correlation coefficient. *: $0.01 < p < 0.05$, **: $0.001 < p < 0.01$, ***: $p < 0.001$. (**C**) The regulatory phosphosites of Erk1/Mapk3 and Erk2/Mapk1. ANOVAs and Dunnett’s post hoc tests were performed on unstimulated (#) or insulin-stimulated (*) phosphoproteome data to compare insulin resistant models to CTRL cells. (**D**) Select phosphosites on canonical insulin signaling proteins that displayed defects in insulin resistance models. (**E**) The distribution of BAS and INS defects among phosphopeptides with defective insulin regulation. For a proportion of defective phosphopeptides we could not significantly attribute defects to BAS or INS (Unassigned). (**F**) Ppp2r5b S13, a phosphosite that displayed emergence in CI, MPQ, and AA. This site was removed from Fig. 2G to improve the heatmap scale. (**G**) GO pathway enrichment on genes containing phosphopeptides that displayed emergent up-regulation (red) or emergent down-regulation (blue) in three or more insulin resistance models. All pathways that were significant after Benjamini-Hochberg p-value adjustment are displayed (only significant for up-regulated phosphopeptides and biological processes), and colour gradients indicate the number of phospho-emergent genes in each pathway.

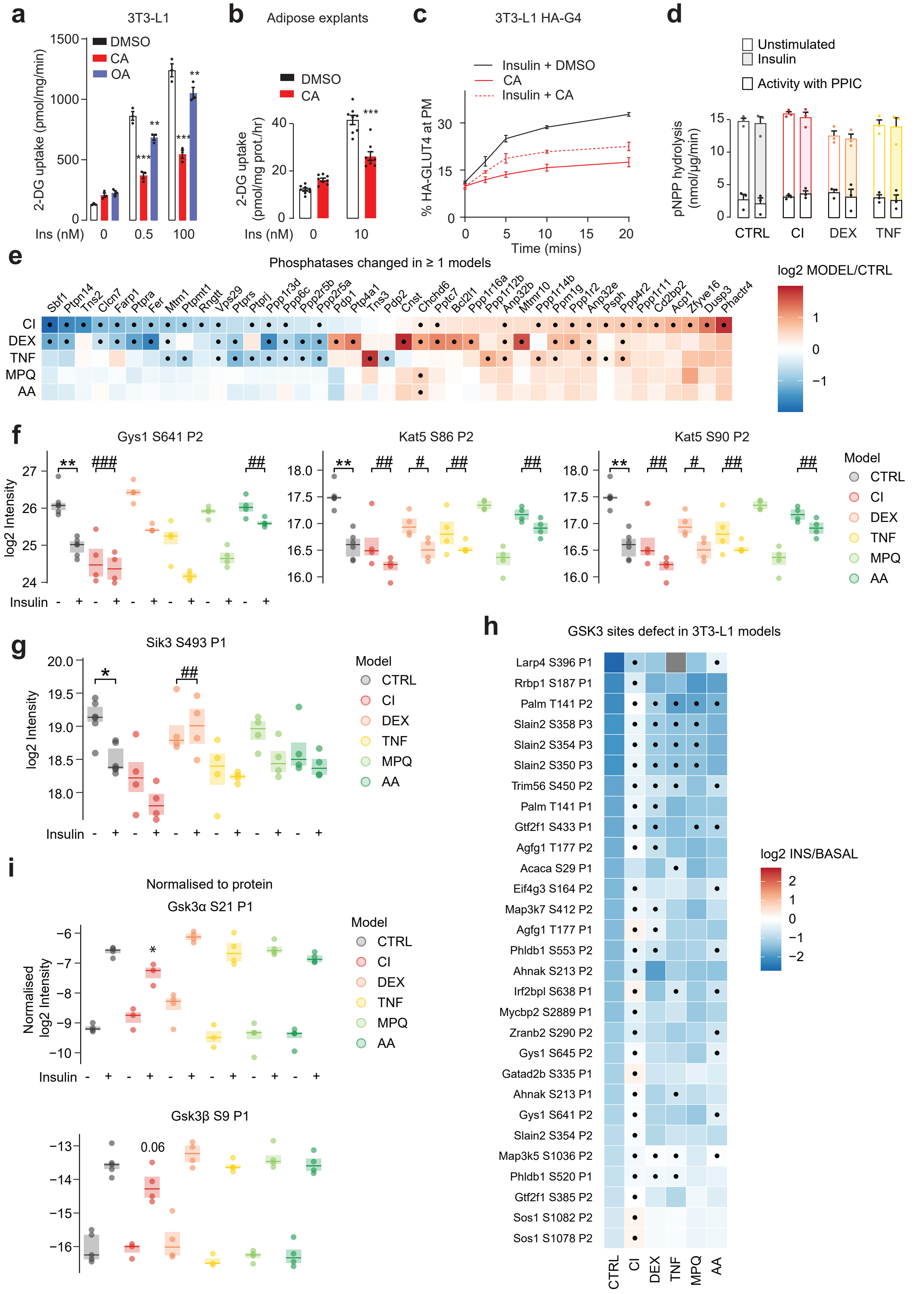
a
3T3-L1
DMSO
CA
OA
2-DG uptake (pmol/mg/min)
Ins (nM)
0
0.5
100
b
Adipose explants
DMSO
CA
2-DG uptake (pmol/mg prot./hr)
Ins (nM)
0
10
c
3T3-L1 HA-G4
Insulin + DMSO
CA
Insulin + CA
% HA-GLUT4 at PM
Time (mins)
d
Unstimulated
Insulin
Activity with PPIC
pNPP hydrolysis (nmol/µg/min)
CTRL
CI
DEX
TNF
e
Phosphatases changed in ≥ 1 models
log2 MODEL/CTRL
Sbf1 Ptpn14 Tns2 Clcn7 Farp1 Ptpra Fer Mtm1 Ptpmt1 Rngtt Vps29 Ptprs Ptprj Ppp1r3d Ppp6c Ppp2r5b Ppp2r5a Pdp1 Ptp4a1 Tns3 Pdp2 Cnst Chchd6 Pptc7 Bcl2l1 Ppp1r16a Ppp1r12b Anp32b Mtmr10 Ppp1r14b Ppm1g Ppp1r2 Anp32e Psph Ppp4r2 Ppp1r11 Cd2bp2 Acp1 Zfyve16 Dusp3 Phactr4
CI
DEX
TNF
MPQ
AA
f
Gys1 S641 P2
Kat5 S86 P2
Kat5 S90 P2
log2 Intensity
Insulin
Model
CTRL
CI
DEX
TNF
MPQ
AA
g
Sik3 S493 P1
log2 Intensity
Insulin
Model
CTRL
CI
DEX
TNF
MPQ
AA
h
GSK3 sites defect in 3T3-L1 models
Larp4 S396 P1
Rrbp1 S187 P1
Palm T141 P2
Slain2 S358 P3
Slain2 S354 P3
Slain2 S350 P3
Trim56 S450 P2
Palm T141 P1
Gtf2f1 S433 P1
Agfg1 T177 P2
Acaca S29 P1
Eif4g3 S164 P2
Map3k7 S412 P2
Agfg1 T177 P1
Phldb1 S553 P2
Ahnak S213 P2
Irf2bpl S638 P1
Mycbp2 S2889 P1
Zranb2 S290 P2
Gys1 S645 P2
Gatad2b S335 P1
Ahnak S213 P1
Gys1 S641 P2
Slain2 S354 P2
Map3k5 S1036 P2
Phldb1 S520 P1
Gtf2f1 S385 P2
Sos1 S1082 P2
Sos1 S1078 P2
CTRL
CI
DEX
TNF
MPQ
AA
log2 INS/BASAL
i
Normalised to protein
Gsk3α S21 P1
Normalised log2 Intensity
Insulin
Model
CTRL
CI
DEX
TNF
MPQ
AA
Gsk3β S9 P1
0.06

**Fig S4: Defective insulin-mediated dephosphorylation in insulin resistance (related to fig 3)** (**A**) $^{3}$H-2DG uptake into 3T3-L1 adipocytes treated with DMSO, calyculin A (CA; 50 nM) or okadaic acid (OA; 1 μM) and insulin at the indicated concentrations for 20 min. Data were analysed by two-way ANOVA corrected for multiple comparisons (Dunnett's test) to compare between DMSO and CA- or OA-treated cells (*). Error bars = S.E.M. n = 3. (**B**) $^{3}$H-2DG uptake into epididymal adipose explants treated with DMSO or CA (100 nM) with insulin at the indicated concentrations for 20 mins. Data were analysed by two-way ANOVA corrected for multiple comparisons (Dunnett's test) to compare between DMSO and CA- or OA-treated cells (*). Error bars = S.E.M. n = 8. (**C**) 3T3-L1 adipocytes overexpressing HA-GLUT4 were treated with insulin (100 nM) and DMSO, CA (50 nM), or insulin and CA. The percentage of HA-GLUT4 at the plasma membrane, as a proportion of total cellular HA-GLUT4, was measured by immunofluorescence at the indicated time points. (**D**) Adipocytes were unstimulated or stimulated with insulin (100 nM) and phosphatase activity was inhibited by phosphatase inhibitor cocktail (PPIC). Total phosphatase activity in 3T3-L1 adipocyte insulin resistant models was measured by hydrolysis of pNPP. Data were analyzed by two-way ANOVA corrected for multiple comparisons (Dunnett's test) to compare between control and insulin resistance models, and unstimulated and insulin-stimulated conditions within models. Error bars = S.E.M. n = 3. (**E**) Phosphatases that were significantly changed in at least one insulin resistance model compared to control cells. Dots indicate significant differences between models and control. (**F**) Select Gsk3 substrates that displayed phosphorylation defects in insulin resistance models. * with bracket: CTRL INS is significantly different to CTRL BAS. # with bracket: Model insulin response is significantly different to CTRL insulin response. (**G**) Sik3 S493, the only insulin-regulated PKA substrate that was defective in insulin resistance models. (**H**) Gsk3 substrates identified through our Gsk3 inhibitor phosphoproteomics that displayed phosphorylation defects in insulin resistance models. Dots indicate that model insulin responses were significantly different to control insulin responses. Missing values are colored grey. (**I**) Intensity of Gsk3α S21 and Gsk3β S9, normalized to the respective total proteins. ANOVAs and Dunnett's post hoc tests were performed on unstimulated (#) or insulin-stimulated (*) phosphoproteome data to compare insulin resistant models to control cells. *: $0.01 < p < 0.05$, **: $0.001 < p < 0.01$, ***: $p < 0.001$.

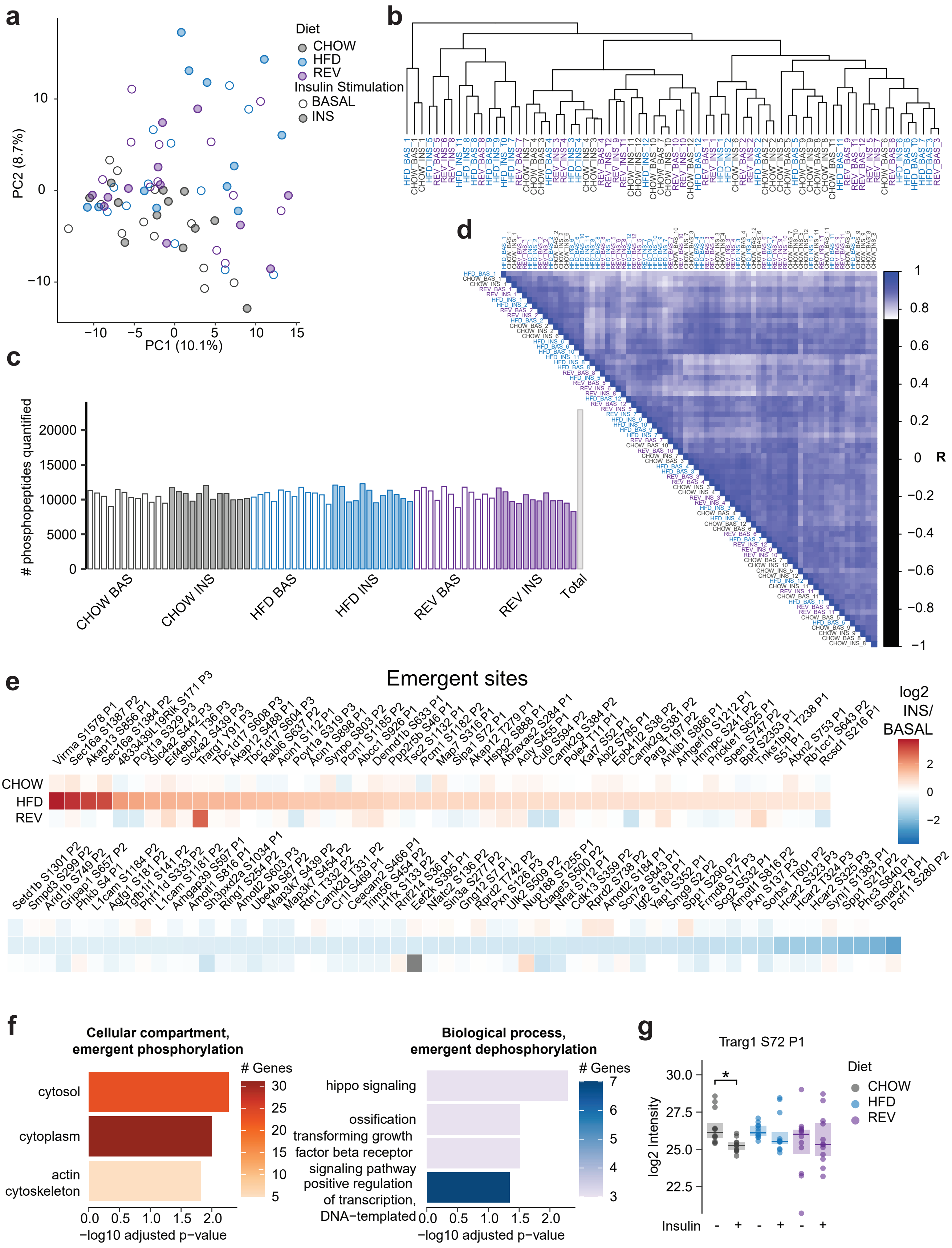
a
Diet
CHOW
HFD
REV
Insulin Stimulation
BASAL
INS
PC2 (8.7%)
PC1 (10.1%)
b
c
### phosphopeptides quantified
CHOW BAS
CHOW INS
HFD BAS
HFD INS
REV BAS
REV INS
Total
d
R
e
Emergent sites
log2 INS/ BASAL
CHOW
HFD
REV
Virma S1578 P1, Sec16a S1387 P2, Akap13 S856 P1, Sec16a S1384 P2, 4833439L19Rik S171 P3, Pcyt1a S329 P3, Slc4a2 S442 P3, Eif4ebp1 T36 P3, Slc4a2 S439 P3, Trarg1 Y91 P3, Tbc1d17 S608 P3, Akap12 S488 P1, Tbc1d17 S604 P3, Rabl6 S637 P2, Acin1 S112 P1, Pcyt1a S319 P3, Acin1 S898 P1, Synpo S803 P2, Pcm1 S1185 P2, Abcc1 S926 P1, Dennd1b S633 P1, Ppp2r5b S46 P1, Tsc2 S1132 P1, Pcm1 S1182 P2, Map7 S316 P1, Sipa1 S72 P1, Akap12 T279 P1, Hspg2 S888 P1, Abraxas2 S284 P1, Acly S455 P1, Cul9 S594 P2, Camk2g S384 P2, Pole4 T11 P1, Kat7 S52 P1, Abl2 S785 P1, Epb41l2 S38 P2, Camk2g S381 P2, Parg T197 P2, Ankib1 S886 P1, Arhgef10 S1212 P1, Hnrnpc S241 P2, Prickle1 S625 P1, Spen S747 P2, Bptf S2353 P1, Tnks1bp1 T238 P1, S51 P1, Atxn2 S753 P1, Rb1cc1 S643 P2, Rcsd1 S216 P1
Setd1b S1301 P2, Smpd3 S299 P2, Arid1b S749 P2, Gripap1 S657 P2, Phkb S4 P1, L1cam S1184 P2, Agfg1 S181 P2, Tgfb1i1 S141 P2, Phf11d S333 P2, L1cam S1181 P2, Arhgap39 S597 P1, Amotl1 S816 P1, Sh3pxd2a S1034 P1, Ring1 S254 P2, Amotl2 S603 P3, Ube4b S87 P2, Map3k7 S439 P2, Map3k7 S454 P2, Rtn1 T332 P2, Camk2d T331 P2, Cr1l S469 P1, Ceacam2 S466 P1, Trim56 S454 P2, H1fx S133 P2, Rnf214 S36 P1, Eef2k S395 P1, Nfatc2 S136 P2, Sin3a S277 P2, Gng12 S7 P1, Rprd2 T742 P2, Pxn S126 P3, Ulk2 S509 P2, Nup188 S1255 P1, Ctage5 S500 P2, Nnat S112 P1, Cdk13 S359 P2, Rprd2 S738 P2, Amotl2 S184 P1, Scn7a S843 P1, Igf2 S183 P1, Yap1 S352 P1, Smg9 S2 P2, Spp1 S290 P2, Frmd8 S17 P3, Scg2 S532 P1, Amotl1 S816 P2, Pxn S137 P3, Sorbs1 T601 P2, Hcar2 S323 P3, Hcar2 T324 P3, Hcar2 S325 P3, Synj1 S1383 P1, Spp1 S212 P2, Phc3 S840 P1, Smad2 T8 P1, Pcf11 S280 P2
f
Cellular compartment, emergent phosphorylation
cytosol
cytoplasm
actin cytoskeleton
### Genes
−log10 adjusted p−value
Biological process, emergent dephosphorylation
hippo signaling
ossification
transforming growth factor beta receptor signaling pathway
positive regulation of transcription, DNA−templated
### Genes
−log10 adjusted p−value
g
Trarg1 S72 P1
log2 Intensity
Diet
CHOW
HFD
REV
Insulin

**Fig S5: Altered insulin signaling in insulin resistant adipose tissue (related to Fig 4)**

(**A**) PCA was performed on the adipose tissue insulin resistance phosphoproteome using the base R "prcomp" function. The first two principal components (PC1 and PC2) are plotted for each replicate, and the percentage contribution of these principal components to total variance is indicated. (**B**) Hierarchical clustering of the phosphoproteome using the base R functions "dist" and "hclust". (**C**) Number of quantified phosphopeptides in each replicate of the phosphoproteome. (**D**) Pearson's correlation between phosphoproteome replicates. (**E**) Emergent phosphopeptides that were regulated by insulin in HFD mice but not in CHOW mice. Missing values are colored gray. (**F**) GO pathway enrichment on genes containing phosphopeptides with emergent up-regulation (red) or down-regulation (blue) in HFD mice. All pathways that were significant after Benjamini-Hochberg p-value adjustment are displayed, and colour gradients indicate the number of phospho-emergent genes in each pathway. (**G**) Trarg1 S72, a Gsk3 substrate that was regulated by insulin in CHOW and defective in HFD mice. * with bracket: INS significantly different to BAS in the indicated condition. *: $0.01 < p < 0.05$, **: $0.001 < p < 0.01$, ***: $p < 0.001$.

**Supplemental data legends**

***Data S1: Proteomic analysis of insulin resistant 3T3-L1 adipocytes***

(Page 1 “quantification”) Normalized LFQ intensities for proteins that passed filtering for further analysis. (Page 2 “analysis”) Statistical analysis of proteome data including models that are changed compared to CTRL.

***Data S2: Proteomic enrichment analysis***

STRING clusters of proteins changed in insulin resistance models. Overrepresentation analysis was performed using KEGG and Reactome pathways on clusters containing 10 or more proteins. (Page 1 “upregulated_clusters”) Clusters containing proteins upregulated in 2 or more models compared to CTRL. (Page 2 “downregulated_clusters”) Clusters containing proteins downregulated in 2 or more models compared to CTRL.

***Data S3: Phosphoproteomic analysis of insulin resistant 3T3-L1 adipocytes***

(Page 1 “quantification”) Normalized LFQ intensities for phosphopeptides that passed filtering for further analysis. (Page 2 “analysis”) Statistical analysis of phosphoproteome data including phosphopeptides that are regulated by insulin in control cells, phosphopeptides with defective responses in insulin resistance models, and phosphopeptides with emergent responses in models.

***Data S4: Canonical insulin signaling proteins***

Gene names for proteins that were considered part of the canonical insulin signaling network.

***Data S5: Phosphoproteomic analysis of 3T3-L1 adipocytes treated with a GSK3 inhibitor***

(Page 1 “quantification”) Normalized LFQ intensities for phosphopeptides that passed filtering for further analysis. (Page 2 “analysis”) Statistical analysis of phosphoproteome data and GSK3 motif analysis. The 290 putative substrate phosphopeptides are identified with the column “Putative substrate”.

***Data S6: Phosphoproteomic analysis of insulin resistant adipose tissue***

(Page 1 “quantification”) Normalized LFQ intensities for phosphopeptides that passed filtering for further analysis. (Page 2 “analysis”) Statistical analysis of phosphoproteome data including phosphopeptides that are regulated by insulin within each diet, phosphopeptides with defective responses in HFD mice, and phosphopeptides with emergent responses in HFD mice.
